# Focusing the GWAS *Lens* on days to flower using latent variable phenotypes derived from global multi-environment trials

**DOI:** 10.1101/2022.03.10.483676

**Authors:** Sandesh Neupane, Derek M Wright, Raul O Martinez, Jakob Butler, James L Weller, Kirstin E Bett

**Author notes:** Author for correspondence: Kirstin E. Bett. These authors contributed equally to this work (Sandesh Neupane, Derek M Wright).

## Abstract

Adaptation constraints within crop species have resulted in limited genetic diversity in some breeding programs and/or areas where new crops have been introduced, *e.g.,* lentil *(Lens culinaris* Medik.) in North America. An improved understanding of the underlying genetics involved in phenology-related traits is valuable knowledge to aid breeders in overcoming limitations associated with unadapted germplasm and expanding their genetic diversity by introducing new, exotic material. We used a large, 18 site-year, multi-environment dataset, phenotyped for phenology-related traits across nine locations and over three years, along with accompanying latent variable phenotypes derived from a photothermal model and principal component analysis (PCA) of days from sowing to flower (DTF) data for a lentil diversity panel (324 accessions) which has also been genotyped with an exome capture array. Genomewide association studies (GWAS) on DTF across multiple environments helped confirm associations with known flowering time genes and identify new quantitative trait loci (QTL), which may contain previously unknown flowering time genes. Additionally, the use of latent variable phenotypes, which can incorporate environmental data such as temperature and photoperiod as both GWAS traits and as covariates, strengthened associations, revealed additional hidden associations, and alluded to potential roles of the associated QTL. Our approach can be replicated with other crop species, and the results from our GWAS serve as a resource for further exploration into the complex nature of phenology-related traits across the major growing environments for cultivated lentil.

## Introduction

The successful introduction of any crop, or new crop varieties, into a new environment is often met with adaptation constraints. Interactions between a genotype and its environment can cause a variety or landrace from one region to perform poorly in another. This can result in genetic bottlenecks as the crop spreads outward from its center of origin, and reluctance from breeders to introduce germplasm from environments dissimilar to their own. These domestication bottlenecks or founder effects have resulted in a loss of genetic diversity in cultivated crops and illustrate the genetic potential which remains underutilized in landraces and crop wild relatives (1–3).

For example, in lentil *(Lens culinaris* Medik.), domestication is estimated to have reduced genetic diversity by 40% (4), and an assessment of genetic diversity among 352 cultivated accessions found narrow genetic variability in what are now the two major production regions for cultivated lentils – South Asia and Western Canada (5). The differences in environmental conditions have driven selection in opposite directions, which in turn inhibits germplasm exchange between the two. South Asian breeding programs are oriented towards early phenology, which is used as an escape mechanism against terminal drought (6), whereas in temperate environments such as Canada, despite a similarly short growing season, later-maturing lines with a longer vegetative growth are preferred to maximize yield. An improved understanding of adaptation or, more specifically, phenology traits such as days from sowing to flower (DTF), an important driving force for adaptation in lentil (7–9), will be necessary to facilitate the introduction of new, exotic germplasm, increasing genetic diversity and enabling breeders to make continued genetic gains.

In general, DTF is controlled by the genotype and its interaction with the controlling environment, which for most crops is predominantly temperature and photoperiod. A pioneer study (8) using six lentil accessions and factorial combinations of temperature and photoperiod in greenhouse experiments, described the rate of progress towards flowering (1/DTF) as a linear function of temperature and photoperiod, a model that has also been used for crops such as pea *(Pisum sativum* L.; (10)), chickpea *(Cicer arietinum* L.; (11, 12)), rice *(Oryza sativa;* (13)), soybean *(Glycine max* L.; (14, 15)), cowpea *(Vigna unguiculata* L., Walp.; (16)), mung bean *(Vigna* spp.; (17)) and faba bean *(Vicia faba* L.; (18, 19)). Later experiments expanded this to a larger, more diverse population of 231 accessions (20) and with field experiments in Syria and Pakistan (21). A more recent study (9) demonstrated the utility of this photothermal model for a lentil diversity panel (LDP), consisting of 324 accessions, phenotyped across 18 site-years representing the three major macro-environments for lentil cultivation *(i.e.,* temperate, South-Asian and Mediterranean environments). This extensive, multi-environment dataset is a useful resource for understanding crop adaptation.

In addition, the recently released annotated lentil genome (22) and single nucleotide polymorphism (SNP) genotyping of the LDP (23) provide an opportunity to combine these data sets and better investigate adaptation in lentil. While a comprehensive review on the genetic control of flowering time has been performed for legumes (24), in lentil most studies are limited to quantitative trait loci (QTL) analyses with bi-parental recombinant inbred line (RIL) populations (25–30), and it remains to be determined if these will be applicable under diverse field conditions, or relevant to a broader set of accessions. Genome-wide association studies (GWAS) have become an important tool for the investigation of complex traits by linking them to specific genomic regions or loci (31), and have been used to identify genes/QTL associated with flowering time in many crop species. Recent GWAS have begun to incorporate environmental factors into the genetic analysis of flowering time through various approaches. These methods can involve GWAS on DTF from differing environments *(e.g.,* Li et al. (32); Li et al. (33); Mao et al. (34)), on differently adapted subgroups *(e.g.,* Huang et al. (35)), on environmental data obtained using passport information *(e.g.,* Navarro et al. (36); Li et al. (33)), or using phenology models to estimate component traits of flowering time (*e.g*., Dingkuhn et al., (37)).

Recently, a new concept called latent phenotyping has arisen in the plant science literature to quantify non-conventional traits (38). This originated from a method called latent space phenotyping (39) developed to detect and quantify response-to-treatment from image data using machine learning and has since been used to quantify whole plant architecture and biomass distribution from LiDAR data of maize hybrids (40). We propose the establishment of a broader and simplified term *latent variable phenotypes* for the plant science community to include any phenotype which is not directly observable, and which may or may not have an easy human interpretation, more similar to the latent variable modeling (41). For example, in the lentil data set from a previous study (9), the principal components (PC1, PC2, PC3) from the PCA on DTF and the coefficients *(a, b, c)* from the photothermal model would all be considered latent variable phenotypes. In addition, the traits derived from the photothermal model, the nominal base temperature (*Tb*), nominal base photoperiod *(Pc),* thermal sum required for flowering *(Tf)* and the photoperiodic sum require for flowering *(Pf)* would also be considered latent variable phenotypes.

The purpose of this study was to evaluate the use of GWAS on a large dataset from multiple environments, with contrasting temperatures and photoperiods, to identify genes/QTL related to phenology and adaptation across the three major macro-environments for cultivated lentil. In addition, by leveraging the use of environmental data through latent variable phenotypes, we improved on the reliability of the GWAS results and were able to associate loci with potential roles based on associated latent variables. This study takes a unique approach, which could be replicated with other crop species, and serves as a resource for further analysis and exploration of genes related to phenology and adaptation.

## Results

### LDP showed high phenotypic variation for DTF across macro-environments

The 324 accessions in the LDP were grown in a total of 18 site-years representing three major macro-environments (9). Average temperatures and photoperiods experienced by the accessions were highly variable across the diverse locations and growing seasons, resulting in significant variation in DTF within and across accessions (**Figure 1**). Temperate environments *(e.g.,* Sutherland, Canada 2017) are characterized by high average temperatures (15.7 °C) and long average photoperiods (16.1 hours), both of which are known to advance flowering in lentil, and therefore showed the lowest DTF values (ranging from 38-62 days) among macro-environments. By contrast, Mediterranean environments (*e.g.*, Metaponto, Italy 2017) had relatively low average temperatures (11.2 °C) and short average photoperiods (10.8 hours), resulting in the highest DTF values recorded (ranging from 97-153 days). South Asian locations (*e.g.*, Jessore, Bangladesh 2017) had short average photoperiods like those described for Mediterranean locations (11 hours) but experienced the highest average temperatures (21.7 °C), resulting in intermediate DTF values (ranging from 45-104).

**Figure 1:**
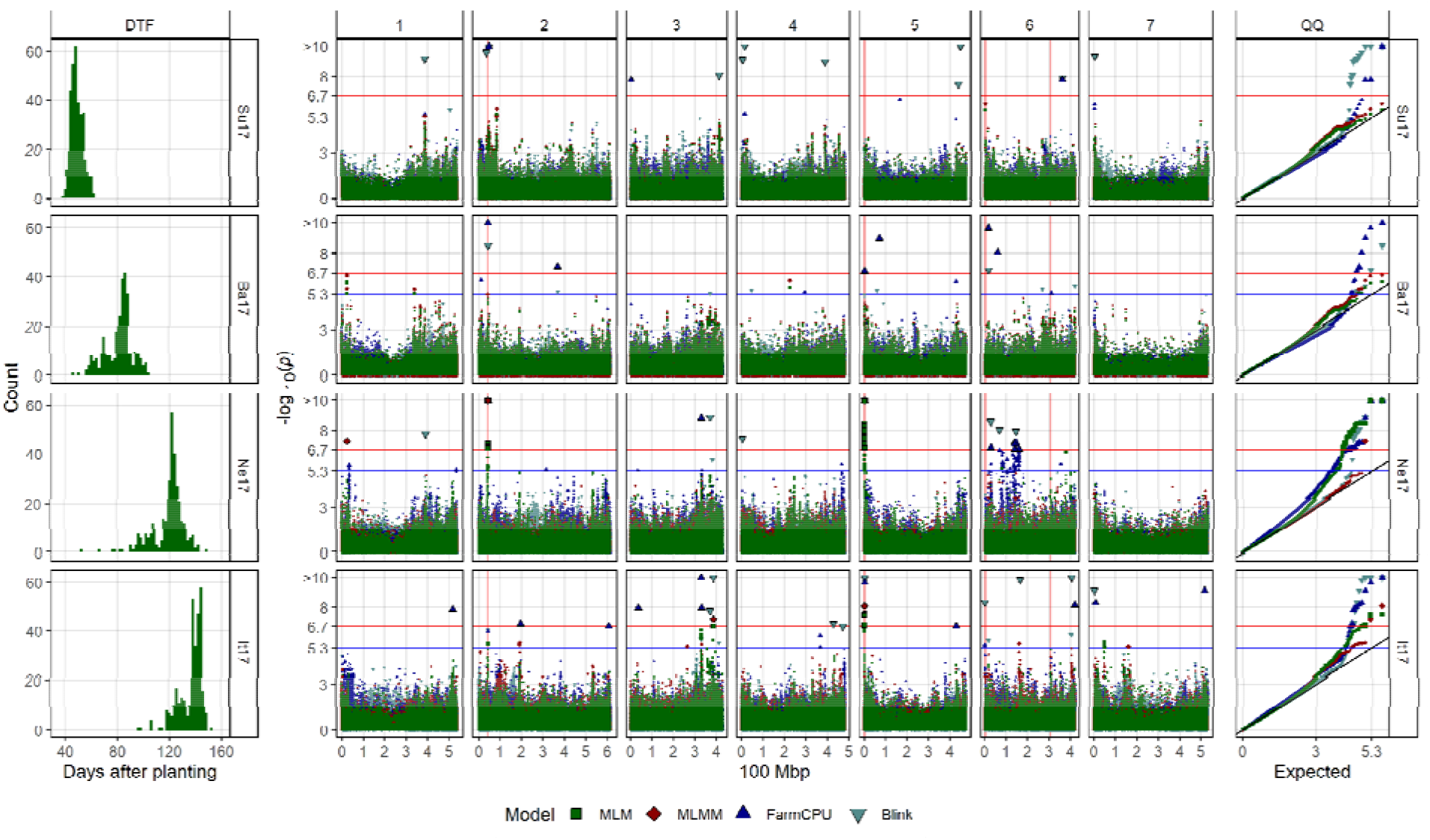
Genome-wide association results for days from sowing to flower (DTF) in Sutherland, Canada 2017 (Su17), Jessore, Bangladesh 2017 (Ba17), Bardiya, Nepal 2017 (Ne17) and Metaponto, Italy 2017 (It17) for a lentil diversity panel. **Left:** Histograms of DTF. **Right:** Manhattan and QQ plots for GWAS results using MLM, MLMM, FarmCPU and Blink models. Vertical lines represent specific base pair locations to facilitate comparisons across traits.

### Latent variable phenotypes improve interpretation of GWAS

To identify genomic regions controlling the variation in phenology observed in the LDP, we performed a series of GWAS analyses. The inclusion of latent variable phenotypes as traits and/or covariates in the GWAS gave important information that alluded to the potential role of the detected QTL within the photoperiod-mediated or temperature-mediated flowering pathways in lentil.

A total of 1086 significantly associated SNPs were detected across all traits and environments (**Figure 2; Supplemental Figure 1; Supplemental Table 1**). Compared to the results from temperate sites, results from Mediterranean and South Asian sites had, in general, a higher number of associations that also showed a higher statistical significance. These differences across macro-environments might be reflecting that any alteration in the flowering pathways would be more evident under the less flowering-inductive conditions of the latter environments than in the highly inductive temperatures and photoperiods that characterize temperate growing seasons. SNPs associated with PC1 were generally also associated with the *b* coefficient of the photothermal model, and a similar co-association was observed between PC2 and the *c* coefficient (**Figure 2; Figure 3**). These results suggests that for the PCA from our previous study (9), the first principal component (PC1) may reflect temperature sensitivity, while the second (PC2) represents photoperiod sensitivity. These two principal components accounted for 68.3% and 14.3% of the variation in DTF across all 18 site-years, respectively (9).

**Figure 2:**
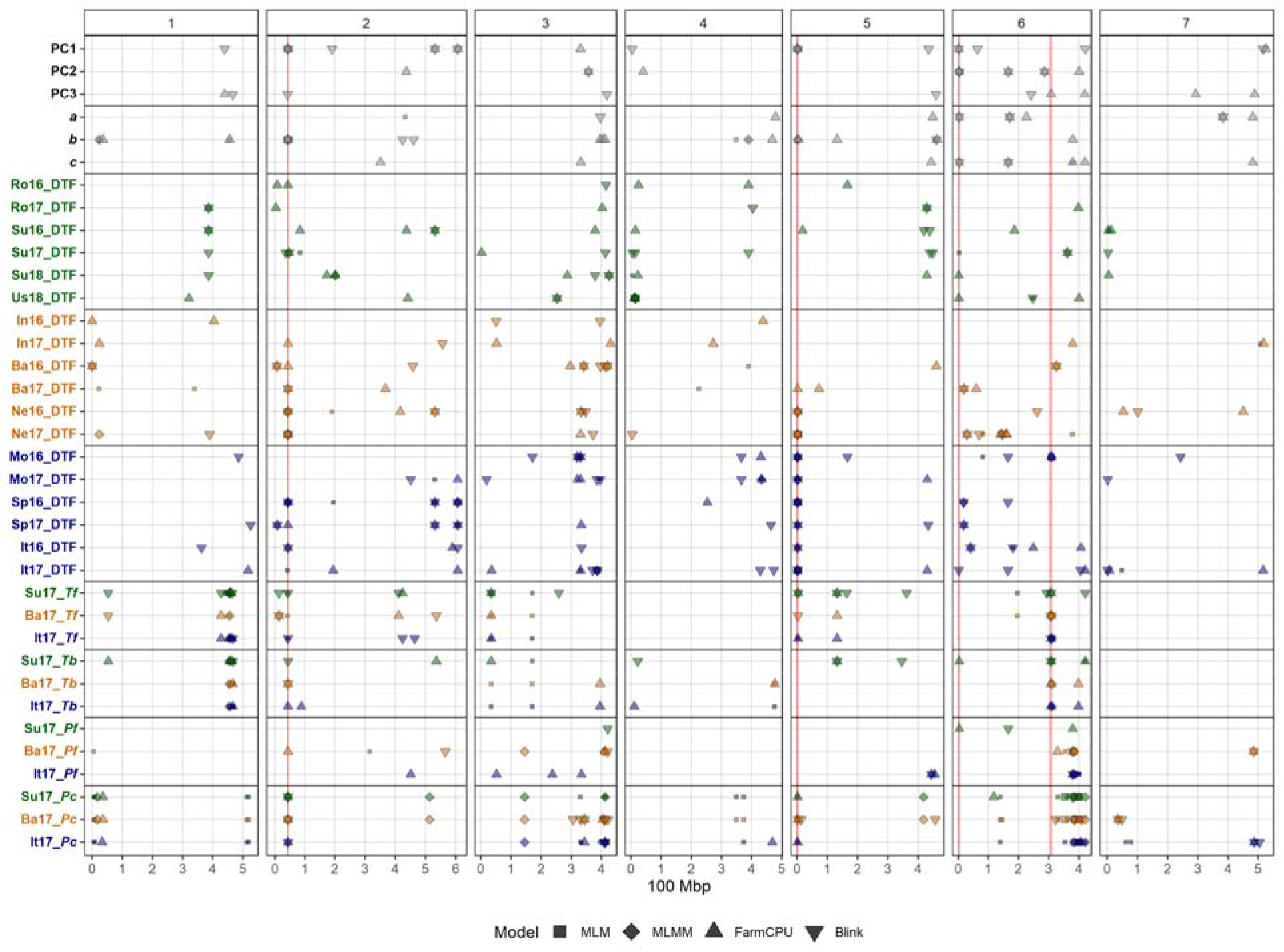
Summary of genome-wide association results using MLM, MLMM, FarmCPU and Blink models on single environment and multi-environment traits related to days from sowing to flower (DTF) in a lentil diversity panel. Larger points represent a significant association (-log10(p) > 6.7) with a trait of interest under one of the GWAS models, while smaller points represent a suggestive association (-log10(p) > 5.3). PC1, PC2, PC3, represent the first three principal components of an analysis of 18 site-years of DTF data (9) in Rosthern, Canada 2016 and 2017 (Ro16 and Ro17), Sutherland, Canada 2016, 2017 and 2018 (Su16, Su17 and Su18), Central Ferry, USA 2018 (Us18), Bhopal, India 2016 and 2017 (In16 and In17), Jessore, Bangladesh 2016 and 2017 (Ba16 and Ba17), Bardiya, Nepal 2016 and 2017 (Ne16 and Ne17), Marchouch, Morocco 2016 and 2017 (Mo16 and Mo17), Cordoba, Spain 2016 and 2017 (Sp16 and Sp17), Metaponto, Italy 2016 and 2017 (It16 and It17). a, b and c are coefficients from a photothermal model developed by Wright et al. (9) and used to calculate the nominal base temperature (Tb), nominal base photoperiod (Pc), thermal sum required for flowering (Tf) and the photoperiod sum required for flowering (Pf). Colors are representative of macroenvironments: Temperate (green), South Asian (orange) Mediterranean (blue) and multi-environment traits (grey). Vertical lines represent specific base pair locations to facilitate comparisons across traits.

**Figure 3:**
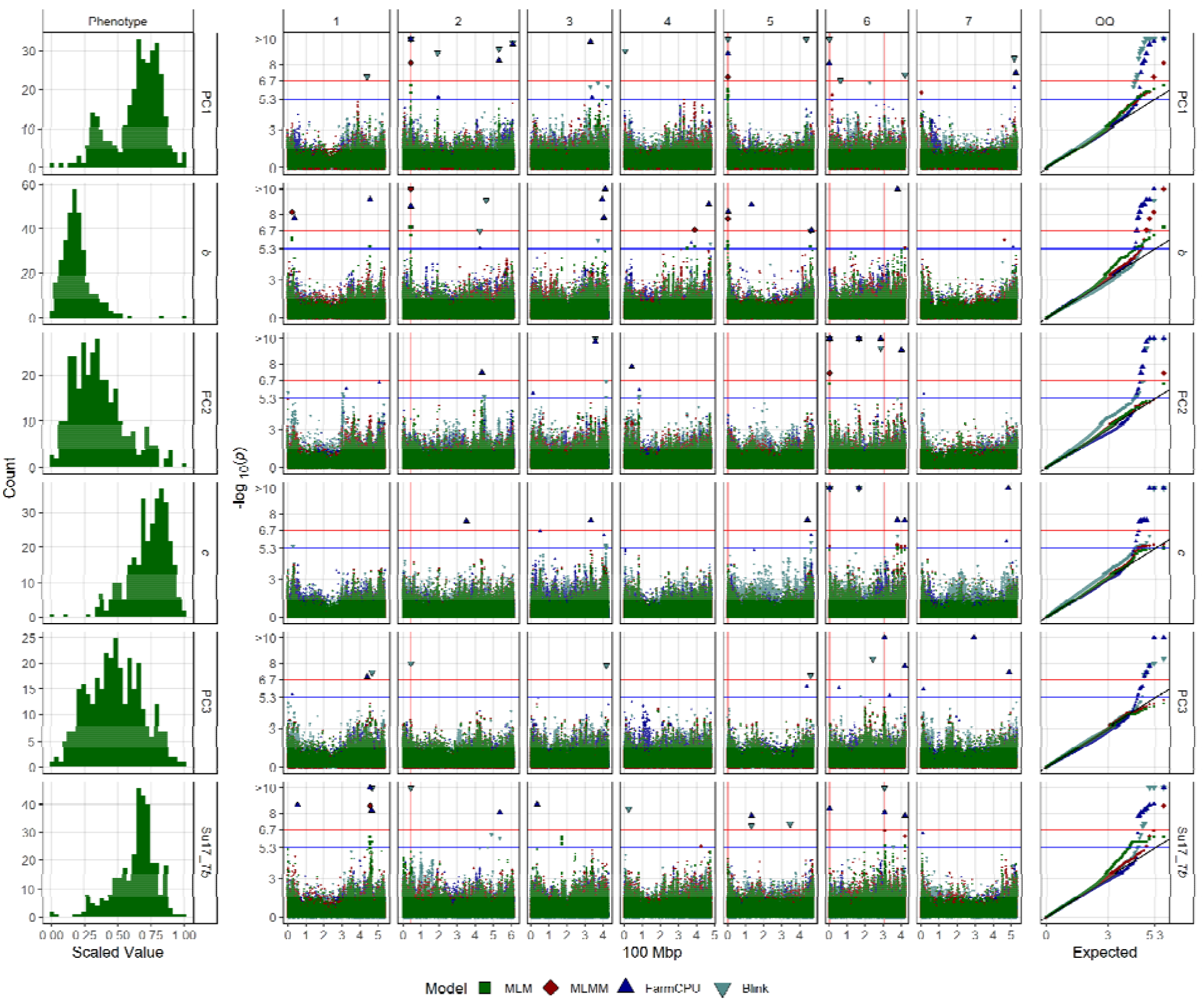
Genome-wide association results for the first three principal components (PC1, PC2 and PC3) the b and c coefficients from the photothermal model developed by Wright et al. (9), and the nominal base temperature (Tb) in Sutherland, Canada 2017 (Su17) for a lentil diversity panel. **Left:** Histograms of each trait, scaled from 0 to 1. **Right:** Manhattan and QQ plots for GWAS results using MLM, MLMM, FarmCPU and Blink models. Vertical lines represent specific base pair locations to facilitate comparisons across traits.

Among the significant SNPs detected by GWAS, one region in chromosome 5, another in chromosome 2 and two in chromosome 6 were repeatedly associated with multiple traits across several environments and/or models (**Figure 2**), and therefore are likely to represent major genes within the molecular pathways governing flowering time in lentil that deserve further investigation.

### Temperature related QTLs identified using latent variable phenotypes

Among the recurring genomic regions, those detected in chromosomes 2 and 5 (with peak SNPs Lcu.2RBY.Chr2p42556949 and Lcu.2RBY.Chr5p1069654 respectively) were strongly associated with the PC1 and the *b* coefficient from the photothermal model (**Figure 3**), suggesting a role in temperature sensitivity. Both QTL showed the largest allelic effect size among all detected by GWAS (**Figure 5a**) and were highly relevant for DTF in both Mediterranean and south Asian locations but not in temperate sites (**Figure 1, Figure 2**). However, when the *c* coefficient was incorporated as a covariate, which should remove photoperiod related associations, they also became significant in temperate locations (**Figure 4a; Supplemental Figure 1**). Similarly, when the *b* coefficient was used as a covariate, which should remove temperature related associations, the associations of these SNPs disappeared in most Mediterranean and South Asian locations (**Figure 4b; Supplemental Figures 1 & 2**), further suggesting a temperature-related role.

**Figure 4:**
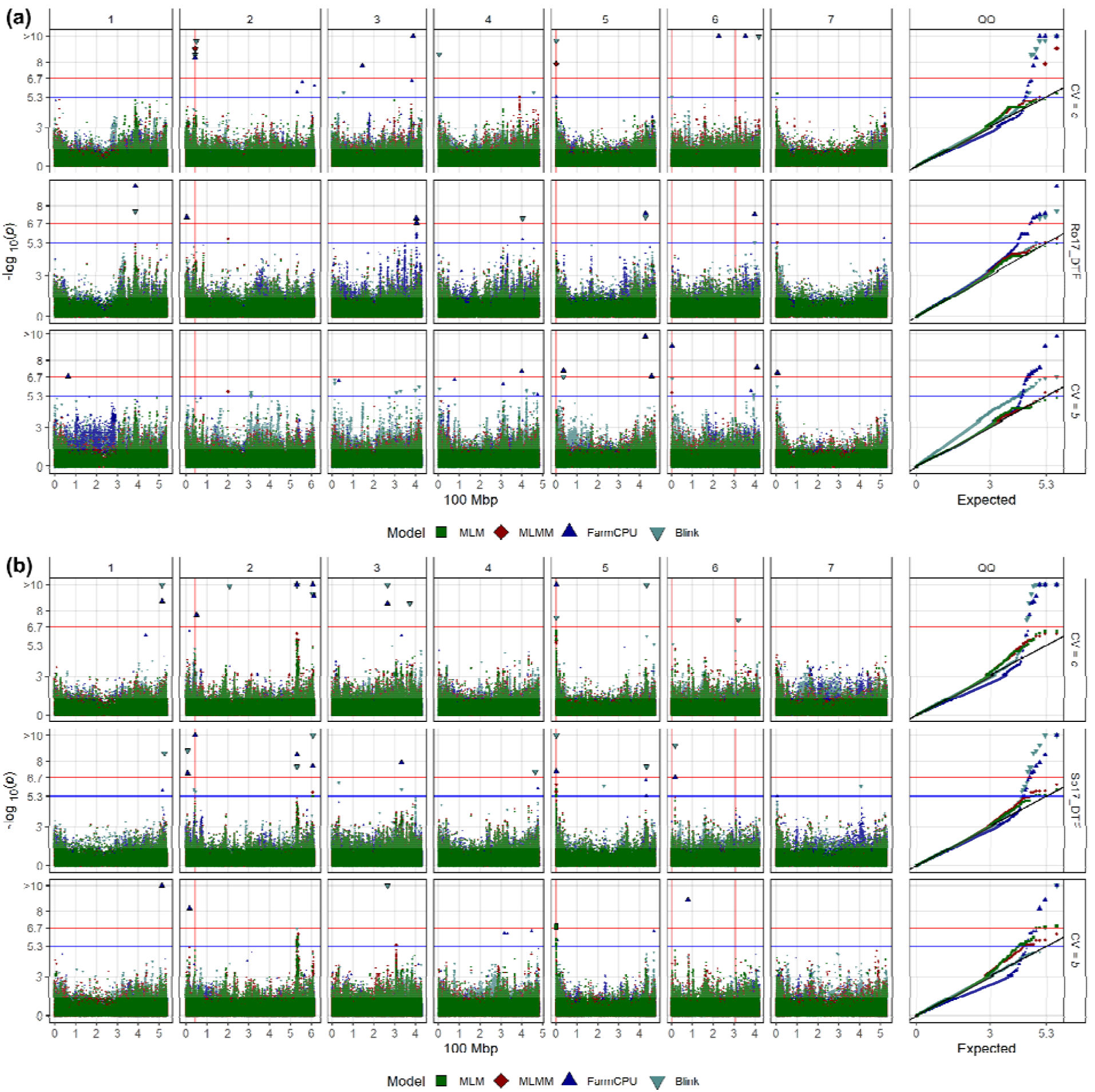
Genome-wide association results for days from sowing to flower (DTF) with and without covariates for a lentil diversity panel. Manhattan and QQ plots for DTF in (a) Rosthern, Canada 2017 (Ro17) and (b) Cordoba, Spain 2017 (Sp17), using MLM, MLMM, FarmCPU and Blink models. The middle panel shows GWAS results without a covariate, while the top and bottom panel show GWAS results using the c and b (temperature and photoperiod) coefficients from the photothermal model developed by Wright et al. (9), respectively. Vertical lines represent specific base pair locations to facilitate comparisons across traits.

**Figure 5:**
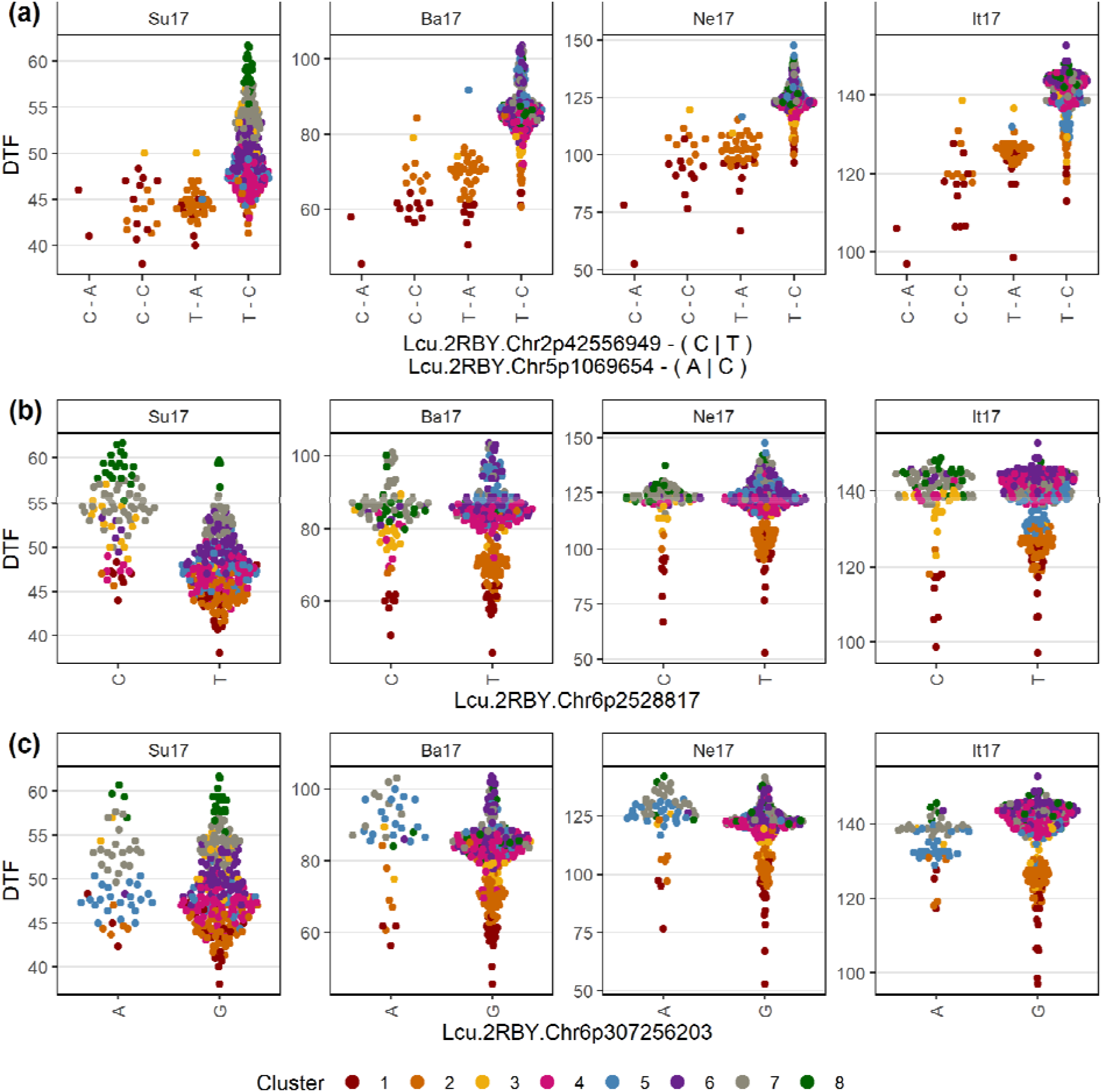
Allelic effects of selected markers on days from sowing to flower (DTF) across contrasting locations in a lentil diversity panel. Sutherland, Canada 2017 (Su17), Jessore, Bangladesh 2017 (Ba17), Bardiya, Nepal 2017 (Ne17) and Metaponto, Italy 2017 (It17). Colors are based on a hierarchical clustering of principal components done by Wright et al. (9) using 18 site-years of DTF data across the three major lentil growing macroenvironments. (a), (b) and (c) correspond to SNPs at the vertical lines on Figures 1, 2, 3 & 4.

The genomic location of these SNPs fell within chromosomal regions lacking any annotated flowering-related genes. A search of potential candidate genes surrounding these SNPs within the window of linkage disequilibrium decay (**Supplemental Figure 4**) revealed several genes with putative links to flowering time (**Supplemental Table 2**, **Supplemental Figure 5**). The most relevant was a *WRKY* gene, belonging to a large family of transcriptional regulators involved in many developmental processes, including flowering time regulation in *Arabidopsis* and rice (42–44). Other candidate genes include a *short-chain dehydrogenase/reductase* similar to the Arabidopsis *SDR6* gene, which has been recently identified as a novel flowering-time gene (45) and three *Ubiquitin-like specific protease (U1pl*). In Arabidopsis, mutations in *Ulp1* orthologs such as *Arabidopsis SUMO Protease1 (ASP1)* and *early in short days4 (ESD4)* are associated with a late and early flowering phenotype, respectively (46, 47). In addition, several other positional candidates were associated with plant response to different stresses (**Supplementary Table 2**). Several studies have shown that, under adverse conditions, plants can accelerate flowering time to secure the production of progeny, and in the last decade this stress-induced flowering has been proposed as an independent flowering pathway (48–51).

A number of SNPs in the central region of chromosome 6 (**Supplemental Figure 3**) were associated with PC1, PC3, DTF in Rosthern17 and with *Tb* and *Tf* across all macroenvironments (**Figure 2**, **Figure 3**). These SNPs define a genomic interval of ~15 Mbp that contains several candidate genes, including a lentil ortholog of the photoperiod related *LWD1* gene (52) and three lentil *FT* orthologs (*LcFTa1, LcFTa2, and LcFTc*), which are well known flowering time genes. In legumes, evidence suggests that *FT* genes acts as integrators of both photoperiod and vernalization (53–57). Interestingly, alleles at Lcu.2RBY.Chr6p307256203 had opposite effects in different environments (**Figure 5c**): in Bangladesh 2017, the minor “A” allele was associated with late flowering, whereas in Italy 2017 it was associated with earliness. Since both environments have similar average photoperiods (11 and 10.8 hours respectively) but different average temperatures (21.7 and 11.2 °C respectively), this observation suggests a possible relation to cold temperatures, which strengthens the potential involvement of the lentil *FT* genes.

### Photoperiod related QTLs identified using latent variable phenotypes

Another QTL in chromosome 6 (Lcu.2RBY.Chr6p2528817) was identified across multiple environments and models. It was strongly associated with the *c* coefficient and PC2 (**Figure 2**, **Figure 3**), indicating that this QTL is likely to participate in the photoperiod response. This QTL was especially relevant for flowering time in Mediterranean environments, and interestingly was also associated with this trait in temperate environments when the *b* coefficient was used as a covariate (**Figure 2**; **Figure 4a**; **Supplemental Figure 1**), further supporting a role in photoperiod response.

This QTL was located at the beginning of chromosome 6 (**Figure 2, Figure 4a**), with a significant SNP in the 5’ UTR of the first of two lentil *FTb* orthologs genes located in tandem at the top of chromosome 6. *FTb* genes are known flowering time genes related to photoperiod response in several legumes, including lentil (24, 53, 56, 58, 59). In all these species, the expression of *FTb* genes show a strong long day requirement, and therefore the effect of a *FTb* allele (i) with a mutation altering their expression pattern would be more relevant in SD environments (Mediterranean, South Asian) or (ii) with a mutation impairing or enhancing its floral inductive capacity would be more evident in locations where photoperiodic requirements are met, such as temperate environments. This can explain the strong allelic effect observed for Lcu.2RBY.Chr6p2528817 in temperate environments compared to Mediterranean and south Asian environments (**Figure 5b**). In winter-sown cropping systems (Mediterranean and South Asia), temperature responsive genes are likely acting first, thus attenuating the effect size of genes related to photoperiod sensitivity.

## Discussion

As in other crops, the spread of lentils from their center of origin in the Fertile Crescent into Europe, Africa, South Asia and eventually to North and South America has been accompanied by a range of physiological adaptations to these new environments that were selected for by early farmers and modern breeders. An appropriate phenology that matches the new environment is an important requirement for adaptation, and in many crops has been achieved through the selection of natural variation in major flowering genes. In several other legume species, most natural variation for flowering time has been relatively well defined genetically, such as the *E* series in soybean, or the four *efl* genes in chickpea (24, 60), and an increasing number of the underlying genes have been molecularly characterized, such as *HR/ELF3a* in pea (61), *Ppd/PHYA* in common bean (62) or the *Ku* locus on narrow-leafed lupin (55). By contrast, knowledge on this matter is scarce in lentil, despite several decades of research into flowering time. To date, only the *ELF3* ortholog *Sn* has been molecularly characterized, with a recessive allele conferring early flowering in a photoperiod-independent manner (29, 61). Other association studies have been published in the last three decades (25–30, 63), but the position of the reported loci cannot be compared due to a lack of common markers between different studies. Identifying the number and (ideally) identity of the genes controlling natural variation in lentil flowering time is of high importance to breeders looking to expand both phenology and genetic diversity, by identifying different combinations of markers/genes which may be absent or underrepresented in their local material.

A standard single environment GWAS approach is clearly insufficient to tease out the full underlying story of the genes controlling lentil flowering time. Here, we have improved the power of our GWAS by using latent variable phenotypes derived from multi-environment trials. These latent variable phenotypes included data derived from a PCA of 18 site-years of DTF data across the three major lentil macro-environments and the incorporation of environmental factors (temperature and photoperiod) with a photothermal model to calculate *b*, *c*, *Tf*, *Pf*, *Tb* and *Pc* (see (9)). To our knowledge, this is the first study to use latent variable phenotypes, which incorporate environmental factors (*b* and *c*), as covariates for DTF in a GWAS, which has helped to confirm associations, allude to their potential roles, and identify others that might have been missed with a single-trait, single-environment approach.

Most of the SNPs detected in our GWAS were locally relevant and likely involved in differential responses to the conditions of one particular site and year. However, four QTL were conspicuously recurrent across different environments and models and are likely to represent variation in important genes within the flowering pathways. Two of these QTL, located in the top and central regions of chromosome 6, co-locate with previously described flowering QTL in lentil. In both cases, members of the lentil *FT* genes have been proposed as strong candidates; a recent study (23) reported the association of the lentil *FTb* genes with flowering time using two different biparental populations, and this year two independent studies supported a role for the *FTa1* gene in lentil flowering (59, 64). Our results serve to further support these conclusions, which in turn increases our confidence in the other associations discovered using the approaches described in this paper.

Two other recurrent QTLs were found, in chromosomes 2 and 5. Their associations with the *b* coefficient and the first principal component from lentil phenological study (9) indicates a temperature-related role. Interestingly, none of the legume flowering genes annotated in the lentil genome fell within the genomic interval defined by LD decay (**Supplemental Figure 4**), indicating that these two loci are likely novel flowering genes. The lack of surrounding known candidates is also indicative of a gap in the general understanding of the flowering network; while the genetic regulation of photoperiod response in plants have been well understood for decades, much less is known about how ambient temperature is integrated into the floral pathways (65).

Despite the strong phenotype of the *sn* allelic variant, which is expected to have a high impact in short day environments such as those described here for the Mediterranean and South Asian locations, we were not able to detect any association between the *Sn* gene with flowering time. A marker designed specifically to detect the causal mutation (a G-to-A substitution in the last nucleotide of the third exon, as described on (61)) revealed that only 7 out of the 324 accessions in the LDP had the early *sn* allele. This illustrates one of the major drawbacks for GWAS, *i.e.,* its limited ability to detect rare or recently generated alleles. The second is the potential for false associations due to population structure. However, our approach using multiple environments, traits and models reduces the possibility of false positives.

It should be emphasized that not all of the variation in DTF can be explained with the 4 major loci discussed here in detail. It is expected that, like other crops, multiple pathways for adaptation to diverse environmental conditions were selected for during lentil geographical expansion. While many of the 1086 significantly associated SNPs identified in this study may turn out to be false positives, further analysis of our results will lead to the identification of more genes/QTL influencing DTF in lentil. A comparison of the cluster groups obtained from a PCA on DTF with the PCA of genetic marker data suggests that there is more to genetic structure than adaptation in lentil, and perhaps a built-in redundancy that allows breeders to maintain some measure of variability (**Supplemental Figure 6**). For example, when considering the two temperature-related QTL on chromosome 2 and 5 (Lcu.2RBY.Chr2p42556949 & Lcu.2RBY.Chr5p1069654), only 2 accessions had both alleles for early flowering, suggesting independent pathways to earliness. Similarly, some of the accessions which contained the late flowering allele for *LcFTb1* (Lcu.2RBY.Chr6p2528817) flowered relatively early, and vice versa, illustrating the potential for other genes/gene combinations to compensate. Interestingly, none of the later flowering accessions with the early flowering allele for *LcFTb1* originated from Canada and represent a large pool of landraces which could be used to expand the genetic diversity within the Canadian germplasm for this particular gene. In addition, our results emphasize the genotype-by-environment interactions which can change how a particular gene/QTL affects DTF, and importantly, the ability/potential to detect associations.

For complex traits such as DTF, which involve a number of interacting mechanisms/pathways influenced by environmental factors, taking the analysis beyond a single environment and adding latent variable phenotypes which incorporate environmental factors proved to be a very effective approach for the genetic dissection of the trait. In just one study we identified 4 genomic regions that could potentially drive lentil adaptation. In other species *(e.g.,* chickpea), the identification of a similar number of genes took decades using more traditional approaches. Two of the detected loci are likely to be novel in legumes, and part of understudied temperature related pathways which are often missed when investigating DTF. The knowledge obtained in this study, and data herein, can/should fuel new research on the topic and serve as a valuable resource for future studies.

## Materials and Methods

### Phenotypic Data

All phenotypic data come from a previous study (9) on a diversity panel consisting of 324 lentil accessions evaluated at 18 site-years across the three major growing environments from 2016 to 2018. Detailed information on the LDP, locations used for trials, and equations for the photothermal model and the traits derived from it *(i.e., a, b, c, Tf, Pf, Tb, Pc)* as well as the PCA results, are defined and described in the same study (9).

### Genotypic Data

Genotyping of all 324 accessions was done using a custom lentil exome capture array as described in previous studies (23, 66). Genotypic data in the form of a high confidence SNP array were accessed from http://knowpulse.usask.ca/AGILE/2. Markers which were not biallelic, had more than 20 accessions with missing data, more than 20% heterozygosity or a minor allele frequency of less than 5% were removed prior to analysis. The remaining 267,846 SNPs, having high coverage across the lentil genome, were used for GWAS.

### Genome-Wide Association Analysis

To thoroughly characterise flowering time across the panel, we used a number of approaches: running GWAS on (1) DTF across varying environments, (2) latent variable phenotypes such as the principal components derived from a PCA on DTF across all environments, and traits derived from a photothermal model which incorporate environmental data, *e.g.*, the temperature and photoperiod coefficients (*b* and *c), Tb, Pc, Tf* and *Pf,* and finally, (3) using latent variable phenotypes, *i.e.*, *b* and *c* coefficients, as covariates in the GWAS to remove confounding associations from the results. These association analyses were performed using different statistical approaches, including Mixed Linear Model (MLM; (67)), Multiple Loci Mixed Model (MLMM; (68)), Fixed and random model Circulating Probability Unification (FarmCPU; (69)) and Bayesian-information and Linkage disequilibrium Iteratively Nested Keyway (BLINK; (70)) models implemented in the Genome Association and Prediction Integrated Tool (GAPIT3; (71)) in R (72). The threshold for significant association was set at 6.7, which is equal to -log_10_(0.05 / 267,846) [i.e. *(p* = 0.05 / no. of markers)], using the Bonferroni correction method to adjust for multiple testing (73). For the quantile-quantile (Q-Q) plots, the observed -log_10_(*p*) values were plotted against the expected -log_10_(*p*) values under the null hypothesis of no association. Further data wrangling and visualization was done using the R packages: “ggbeeswarm” (74), “ggpubr” (75), “ggtext” (76), “glue” (77), “magick” (78), “plyr” (79) and “tidyverse” (80).

### Candidate Gene Identification

Regions of the genome that were significantly associated with DTF and other latent variable phenotypes were identified by filtering the significant markers from the GWAS results. The physical positions of these markers were examined in the annotated lentil genome (22) on JBrowse (https://knowpulse.usask.ca/jbrowse/Lens-culinaris/2). Known legume flowering time genes have been annotated on the lentil genome and are found in the curated genes track of this JBrowse. After identifying candidate genes or genomic regions harbouring significant markers, the allelic proportions at the most significant marker were determined.

## Supporting Information

**Supplemental Figure 1**: Summary of genome-wide association results using MLM, MLMM, FarmCPU and Blink models on days from sowing to flower (DTF) in a lentil diversity panel using no covariates or with the either the *b* or c coefficient as a covariate. Larger points represent a significant association (-log_10_(*p*) > 6.7) with a trait of interest under one of the GWAS models, while smaller points represent a suggestive association (-log_10_(*p*) > 5.3). Rosthern, Canada 2016 and 2017 (Ro16 and Ro17), Sutherland, Canada 2016, 2017 and 2018 (Su16, Su17 and Su18), Central Ferry, USA 2018 (Us18), Bhopal, India 2016 and 2017 (In16 and In17), Jessore, Bangladesh 2016 and 2017 (Ba16 and Ba17), Bardiya, Nepal 2016 and 2017 (Ne16 and Ne17), Marchouch, Morocco 2016 and 2017 (Mo16 and Mo17), Cordoba, Spain 2016 and 2017 (Sp16 and Sp17), Metaponto, Italy 2016 and 2017 (it16 and It17). *b* and *c* are coefficients derived from a photothermal model developed by Wright et al. (9). Colors are representative of macroenvironments: temperate (green), South Asian (orange) and Mediterranean (blue). Vertical lines represent specific base pair locations to facilitate comparisons across traits.

**Supplemental Figure 2**: Regional genome-wide association results from 35 Mbp to 50 Mbp on chromosome 2 for selected traits with a lentil diversity panel using MLM, MLMM, FarmCPU and Blink models. Traits include days from sowing to flower (DTF) in Sutherland, Canada 2017 (Su17), with and without the *c* coefficient used as a covariate (CV=*c*), DTF in Jessore, Bangladesh 2017 (Ba17), DTF in Bardiya, Nepal 2017 (Ne17), DTF in Metaponto, Italy 2017 (It17), the first principal component (PC1) from a principal component analysis of an analysis of 18 site-years of DTF data, and the *b* coefficient derived from a photothermal model done by Wright et al. (9). The vertical line represents a specific base pair location to facilitate comparisons across traits.

**Supplemental Figure 3**: Regional genome-wide association results from 300 Mbp to 315 Mbp on chromosome 6 for selected traits with a lentil diversity panel using MLM, MLMM, FarmCPU and Blink models. Traits include days from sowing to flower (DTF) in Marchouch, Morocco 2016 (Mo16), thermal sum required for flowering (*Tf*) in Sutherland, Canada 2017 (Su17) and Metaponto, Italy 2017 (It17), nominal base temperature (*Tb*) in Su17 and It17, and the third principal component (PC3) from a principal component analysis of an analysis of 18 site-years of DTF data from Wright et al. (9). Vertical lines represent the locations of selected flowering time genes within the associated QTL.

**Supplemental Figure 4**: Linkage disequilibrium decay across the 7 chromosomes in the lentil genome. (a) Linkage disequilibrium decay for all marker combinations. (b) Linkage disequilibrium decay for marker combinations within a 1 Mbp distance. For each chromosome, 1000 SNP were randomly selected for pairwise LD calculations. Shaded lines represent the moving average of 100 pair-wise marker comparisons. Solid line represents a loess regression used to determine the value (vertical dashed line) in which R^2^ reaches the 0.2 threshold (blue dashed line). Red dashed lines represent the average R^2^ for each chromosome.

**Supplemental Figure 5**: Diagram of a 290 kbp region of lentil chromosome 2 (A) and a 215 kbp region of lentil chromosome 5 (B) containing the relevant SNPs Lcu.2RBY.Chr2p42543877, Lcu.2RBY.Chr2p42556949 and Lcu.2RBY.Chr5p1063138 (highlighted in red). The syntenic regions in chickpea and Medicago genomes are also shown for comparison. The length of the relevant interval for each chromosome was calculated according to the SNP position ± chromosome-specific linkage disequilibrium decay. Numbers over or besides each gene correspond to those shown in Supplemental Table 2.

**Supplemental Figure 6**: Pair-wise plots of a principal component analysis of genetic marker data from a lentil diversity panel. Colors are based on a hierarchical clustering of principal components done by Wright et al. (9) using 18 site-years of days from sowing to flower data across the three major lentil growing macroenvironments.

**Supplemental Table 1**: GWAS results for SNPs significantly associated with the traits of interest used in this study. Traits include days from sowing to flower (DTF), the first three principal components from a principal component analysis (PCA) of the DTF data (PC1, PC2, PC3), the *a*, *b* and *c* coefficients from a photothermal model (PTModel), the nominal base temperature (*Tb*), nominal base photoperiod (*Pc*), thermal sum required for flowering (*Tf*) and the photoperiod sum required for flowering (*Pf*). Rosthern, Canada 2016 and 2017 (Ro16 and Ro17), Sutherland, Canada 2016, 2017 and 2018 (Su16, Su17 and Su18), Central Ferry, USA 2018 (Us18), Bhopal, India 2016 and 2017 (In16 and In17), Jessore, Bangladesh 2016 and 2017 (Ba16 and Ba17), Bardiya, Nepal 2016 and 2017 (Ne16 and Ne17), Marchouch, Morocco 2016 and 2017 (Mo16 and Mo17), Cordoba, Spain 2016 and 2017 (Sp16 and Sp17), Metaponto, Italy 2016 and 2017 (It16 and It17). For further details see Wright et al. (9). Traits run with the *b* or *c* coefficients as a covariate are indicated with the “-b” and “-c” suffix in the trait column. https://github.com/derekmichaelwright/AGILE_LDP_GWAS_Phenology/blob/master/Supplemental_Table_01.csv

**Supplemental Table 2**: List of genes in the regions associated with flowering time in lentils chromosomes 2 and 5, and the syntenic regions of chickpea *(Cicer arietinum)* and *Medicago truncatula*. https://github.com/derekmichaelwright/AGILE_LDP_GWAS_Phenology/blob/master/Supplemental_Table_02.xlsx

## Acknowledgements

This research was conducted as part of the “Application of Genomics to Innovation in the Lentil Economy (AGILE)”, a project funded by Genome Canada and managed by Genome Prairie. We are grateful for the matching financial support from the Saskatchewan Pulse Growers, Western Grains Research Foundation, the Government of Saskatchewan, and the University of Saskatchewan. We acknowledge the technical assistance of the bioinformatics, field and molecular lab staff of the Pulse Crop Breeding and Genetics group at the University of Saskatchewan.

## Author Contributions

Conceptualization: Kirstin E Bett, Sandesh Neupane, Derek M Wright

Data Wrangling: Derek M Wright, Sandesh Neupane, Kirstin E Bett

Formal Analysis: Derek M Wright

Funding Acquisition: Kirstin E. Bett

Methodology: Kirstin E Bett, Derek M Wright, Sandesh Neupane, Raul O Martinez, Jakob Butler, James L Weller

Writing – Original Draft: Sandesh Neupane, Derek M Wright, Raul O Martinez

Writing – Review and Editing: Derek M Wright, Sandesh Neupane, Kirstin E Bett, Raul O Martinez, Jakob Butler, James L Weller

## Data Availability

The data supporting this study are available through http://knowpulse.usask.ca/AGILE/2 or from the authors upon request. The source code for all data analyses is available at: https://derekmichaelwright.github.io/AGILE_LDP_GWAS_Phenology/GWAS_Phenology_Vignette.html

