## Supplementary figures and images for "Focusing the GWAS *Lens* on days to flower using latent variable phenotypes derived from global multi-environment trials"

### Supplemental Figure 1

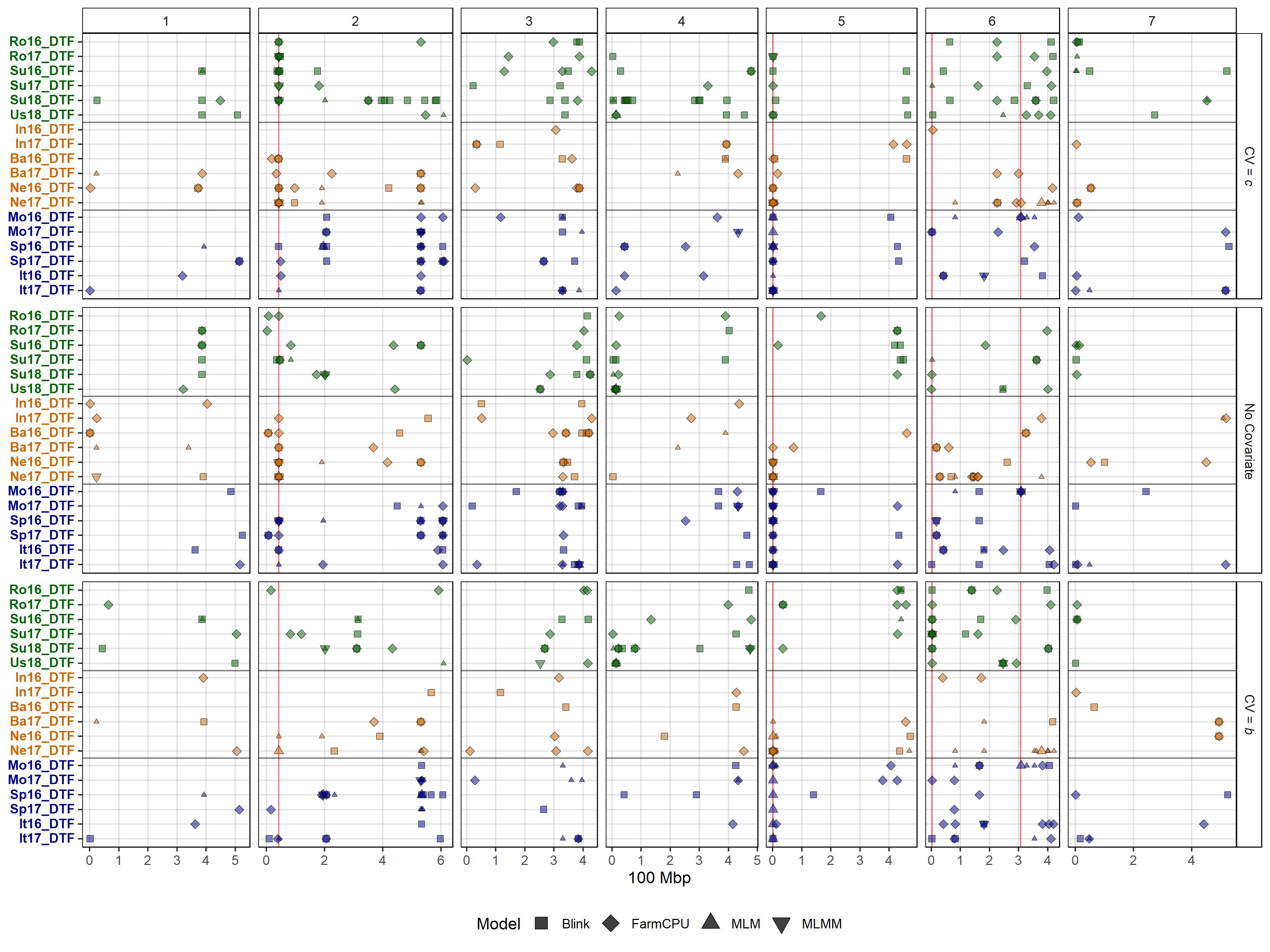

### Supplemental Figure 2

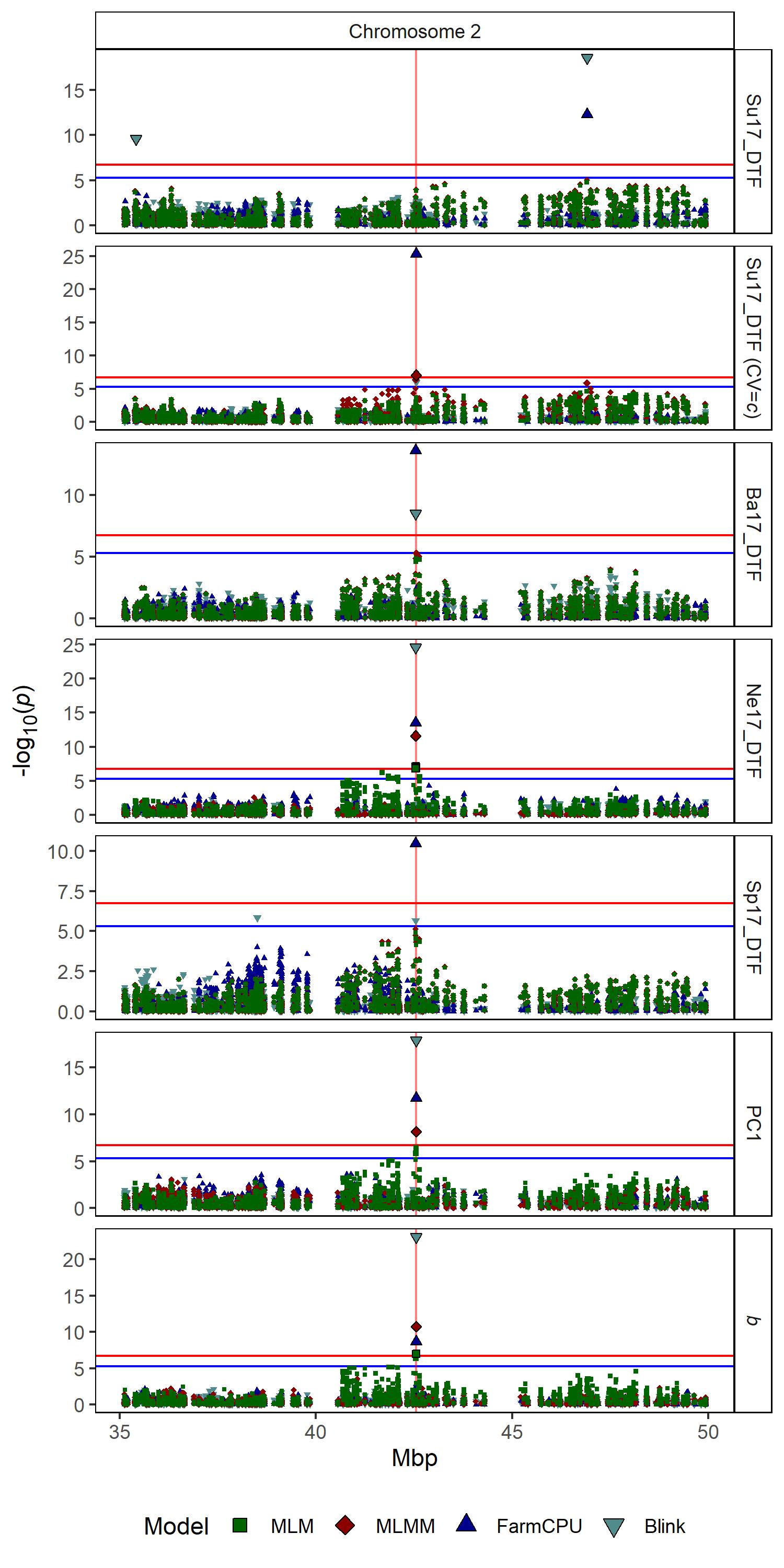

### Supplemental Figure 3

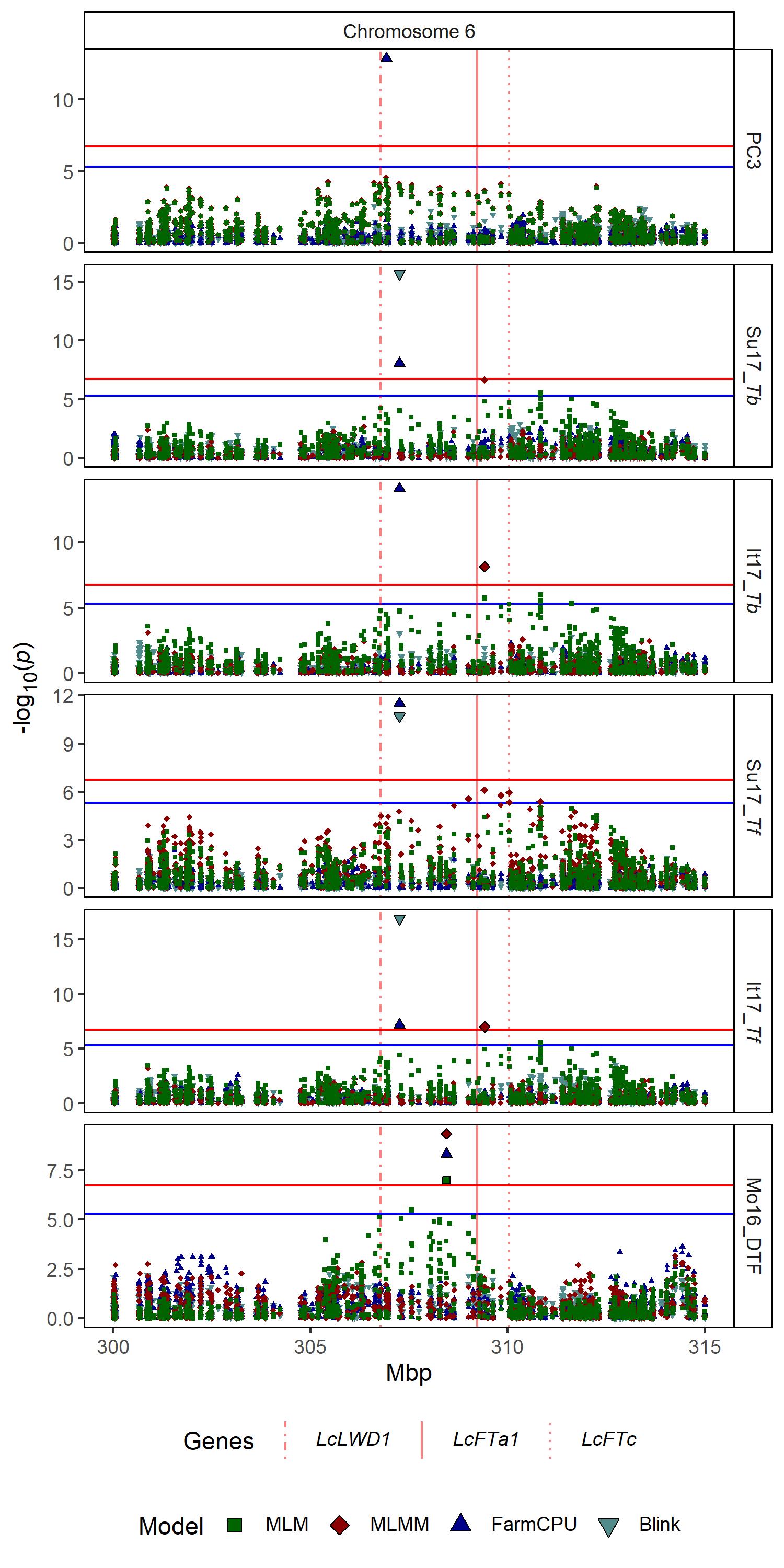

### Supplemental Figure 4

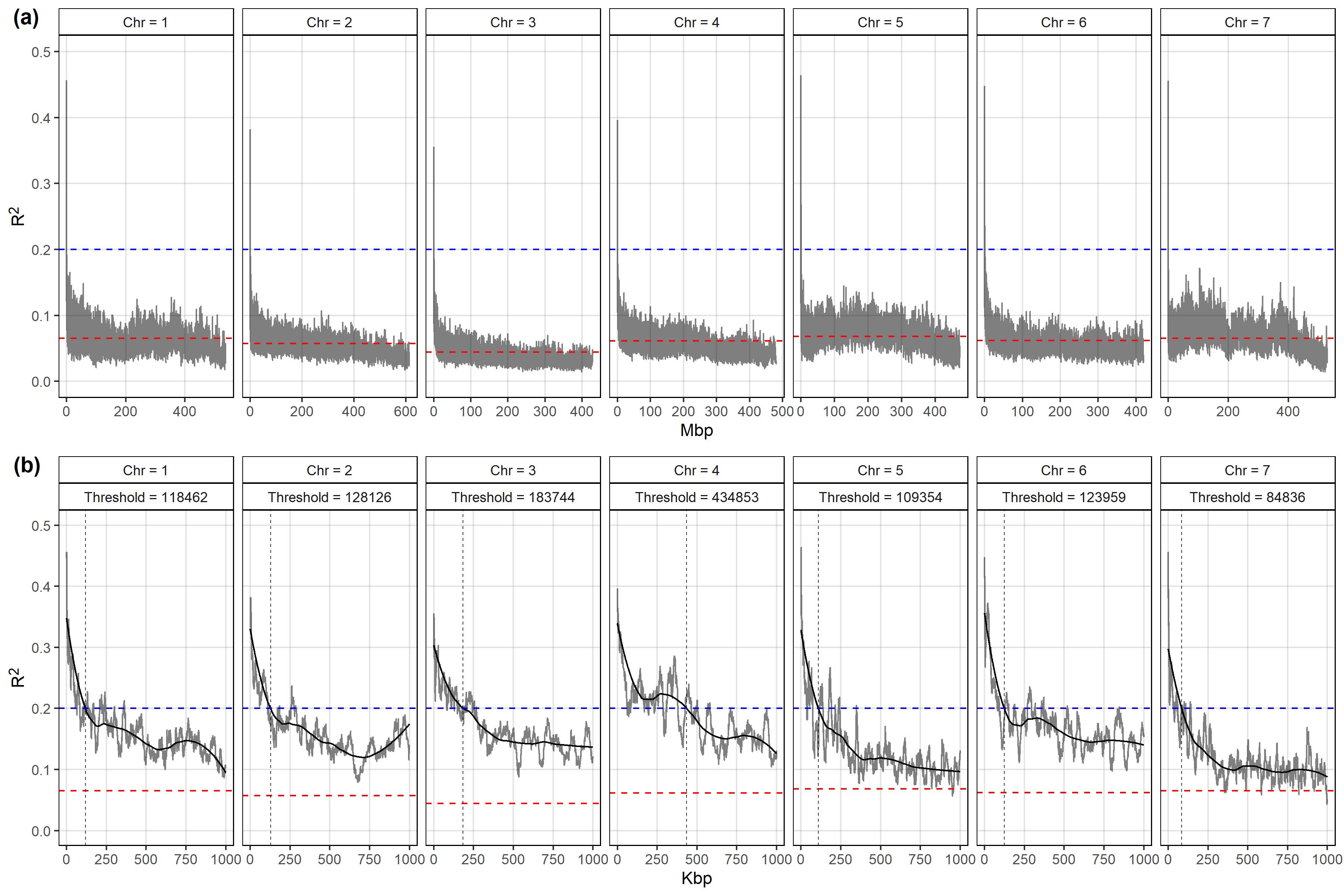

### Supplemental Figure 5

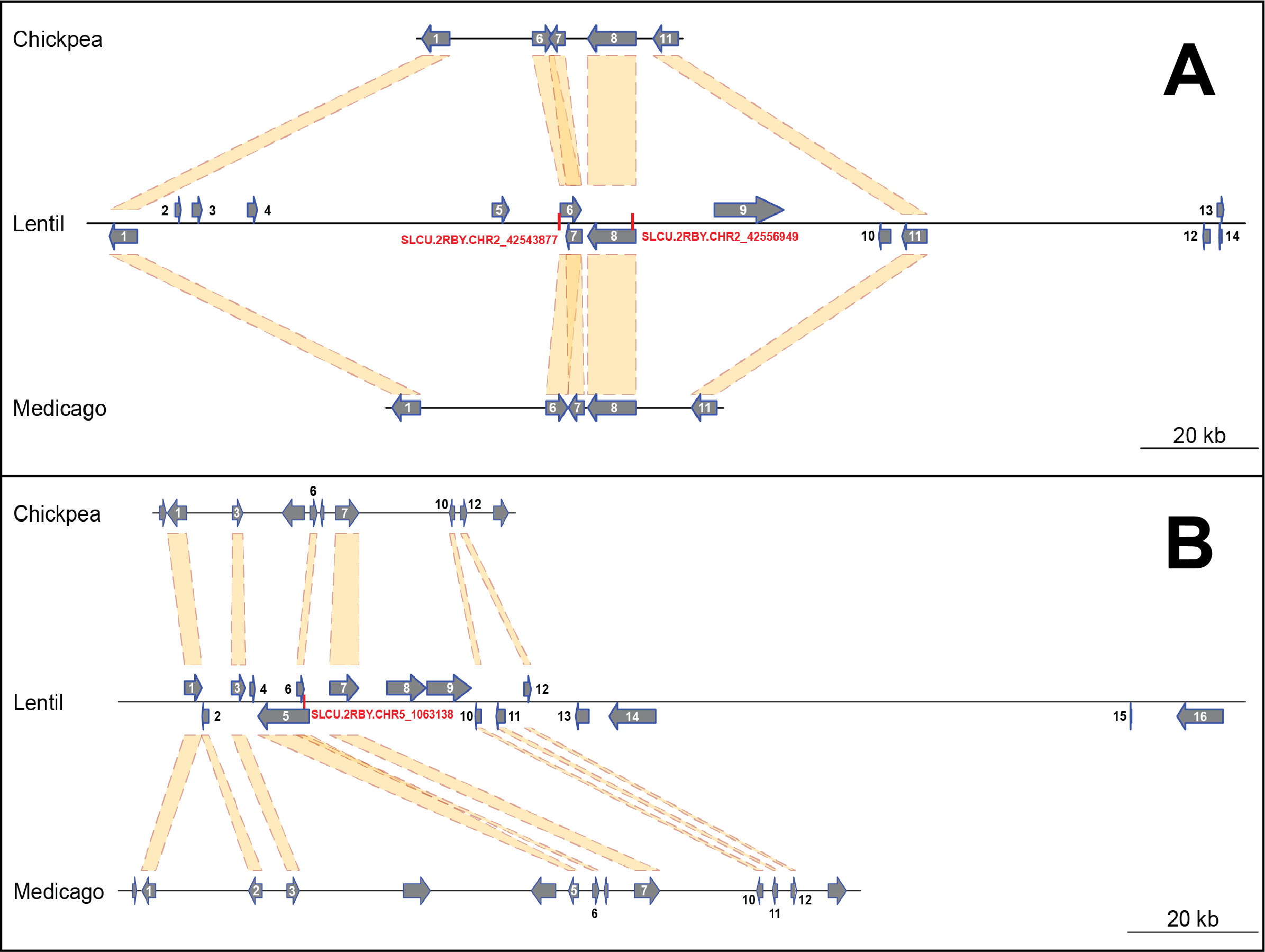

### Supplemental Figure 6

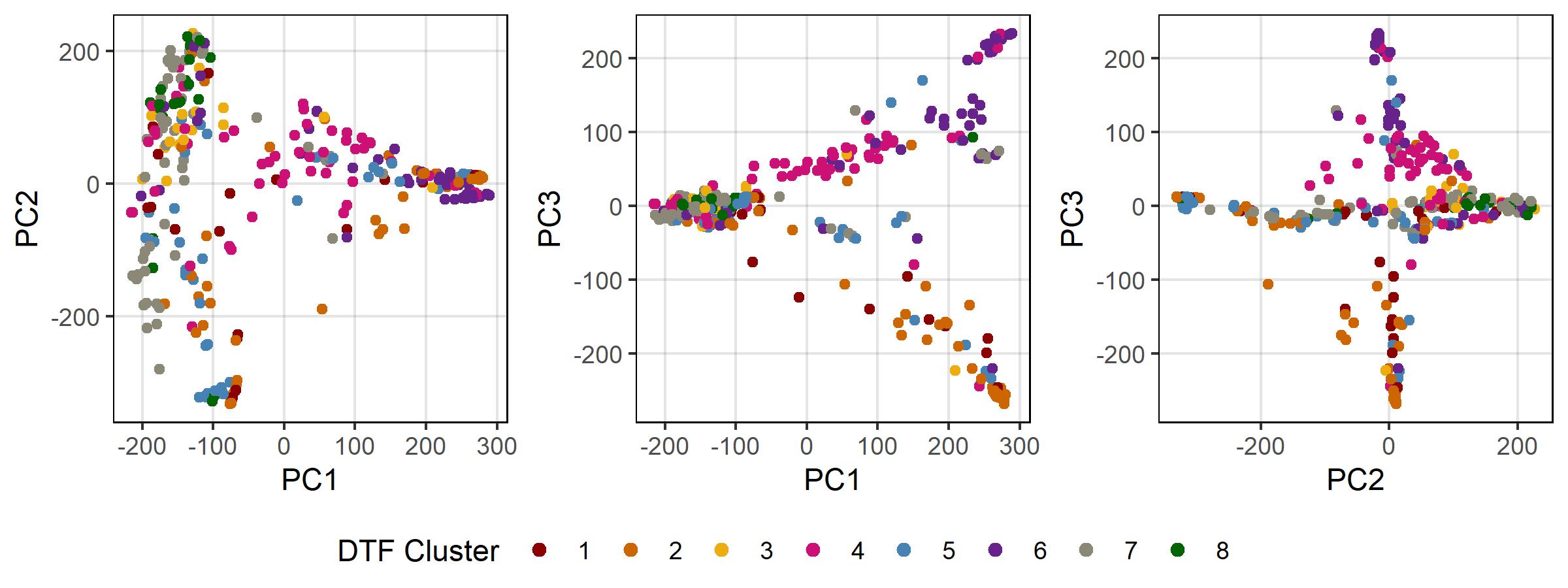
